# Small amounts of misassembly can have disproportionate effects on pangenome-based metagenomic analyses

**DOI:** 10.1101/2024.10.11.617902

**Authors:** Stephanie N. Majernik, Larry Beaver, Patrick H. Bradley

## Abstract

Individual genes from microbiomes can drive host-level phenotypes. To help identify such candidate genes, several recent tools estimate microbial gene copy numbers directly from metagenomes. These tools rely on alignments to pangenomes, which in turn are derived from the set of all individual genomes from one species. While large-scale metagenomic assembly efforts have made pangenome estimates more complete, mixed communities can also introduce contamination into assemblies, and it is unknown how robust pangenome-based metagenomic analyses are to these errors. To gain insight into this problem, we re-analyzed a case-control study of the gut microbiome in cirrhosis, focusing on commensal Clostridia previously implicated in this disease. We tested for differentially prevalent genes in the *Lachnospiraceae*, then investigated which were likely to be contaminants using sequence similarity searches. Out of 86 differentially prevalent genes, we found that 33 (38%) were probably contaminants originating in taxa such as *Veillonella* and *Haemophilus*, unrelated genera that were independently correlated with disease status. Our results demonstrate that even small amounts of contamination in metagenome assemblies, below typical quality thresholds, can threaten to overwhelm gene-level metagenomic analyses. However, we also show that such contaminants can be accurately identified using a method based on gene-to-species correlation. After removing these contaminants, we observe that several flagellar motility gene clusters in the *Lachnospira eligens* pangenome are associated with cirrhosis status. We have integrated our analyses into an analysis and visualization pipeline, PanSweep, that can automatically identify cases where pangenome contamination may bias the results of gene-resolved analyses.

**Importance:** Metagenome-assembled genomes, or MAGs, can be constructed without pure cultures of microbes. Large scale efforts to build MAGs have yielded more complete pangenomes (i.e., sets of all genes found in one species). Pangenomes allow us to measure strain variation in gene content, which can strongly affect phenotype. However, because MAGs come from mixed communities, they can contaminate pangenomes with unrelated DNA, and how much this impacts downstream analyses has not been studied. Using a metagenomic study of gut microbes in cirrhosis as our test case, we investigate how contamination affects analyses of microbial gene content. Surprisingly, even small, typical amounts of MAG contamination (<5%) result in disproportionately high levels of false positive associations (38%). Fortunately, we show that most contaminants can be automatically flagged, and provide a simple method for doing so. Furthermore, applying this method reveals a new association between cirrhosis and gut microbial motility.

## Introduction

The gain or loss of individual genetic elements can drastically change microbial phenotypes, such as the pathogenicity island *cag* in *Helicobacter pylori* (1), the phage-borne virulence factor *stx* in *Shigella dysenteriae* and *Escherichia coli* (2), or the transposon-borne *Bacteroides fragilis* toxin BFT (3). The same applies to commensals: for example, specific strains of commensal *Bacteroides fragilis* encode genes for a capsular polysaccharide, PSA, which in turn evokes an anti-inflammatory response (4). This phenomenon has motivated the development of tools to measure strain-level variation in gene content. Several recent tools can estimate gene content directly from microbial communities, including MIDAS (5, 6), PanPhlAN (7, 8), and StrainPanDA (9). These methods involve aligning reads to species-specific catalogs of genes, i.e., pangenomes, and are therefore faster and require less coverage than metagenomic assembly.

A major drawback, however, is that these tools require many high-quality genomes to estimate pangenomes. While we have thousands of isolate genomes for some species (like *E. coli*), others may have few to none. One solution to this problem is to integrate metagenome-assembled genomes, or MAGs, which do not require pure culture and can be obtained at scale from existing data. Indeed, the construction of large MAG collections for human (10), mouse (11), and environmental (12) microbiomes has enabled much more complete pangenome estimates for a wide range of microbes, including those with few or no cultured representatives. However, genomes may also erroneously include sequences from unrelated taxa (“contamination”), which can then be propagated to pangenomes. Even isolate genomes are not immune to contamination (13), but because they are made by assembling and binning shotgun reads from mixed communities, MAGs are typically more likely to be contaminated by contigs from an unrelated species (14).

Tools such as CheckM (15) and GUNC (16) can be used to identify heavily contaminated MAGs and to exclude them from pangenomes. However, a typical threshold for a “high-quality draft” MAG might still allow up to 5% contamination (17). Additionally, current tools may fail to detect contaminant contigs when they do not include any highly conserved marker genes (14). Thus, a small number of contaminant genes may be introduced into the pangenome catalogue (Figure 1A). This problem has been previously identified (14, 18), but has additional potential consequences in the context of gene-level metagenomic analyses. When reads are aligned to these contaminating genes, they will be attributed to the wrong species. Furthermore, if the source of the contamination is itself changing in abundance in cases vs. controls, the contaminating gene will have the same pattern and would therefore be falsely identified as significantly different (Figure 1B).

**Figure 1.**
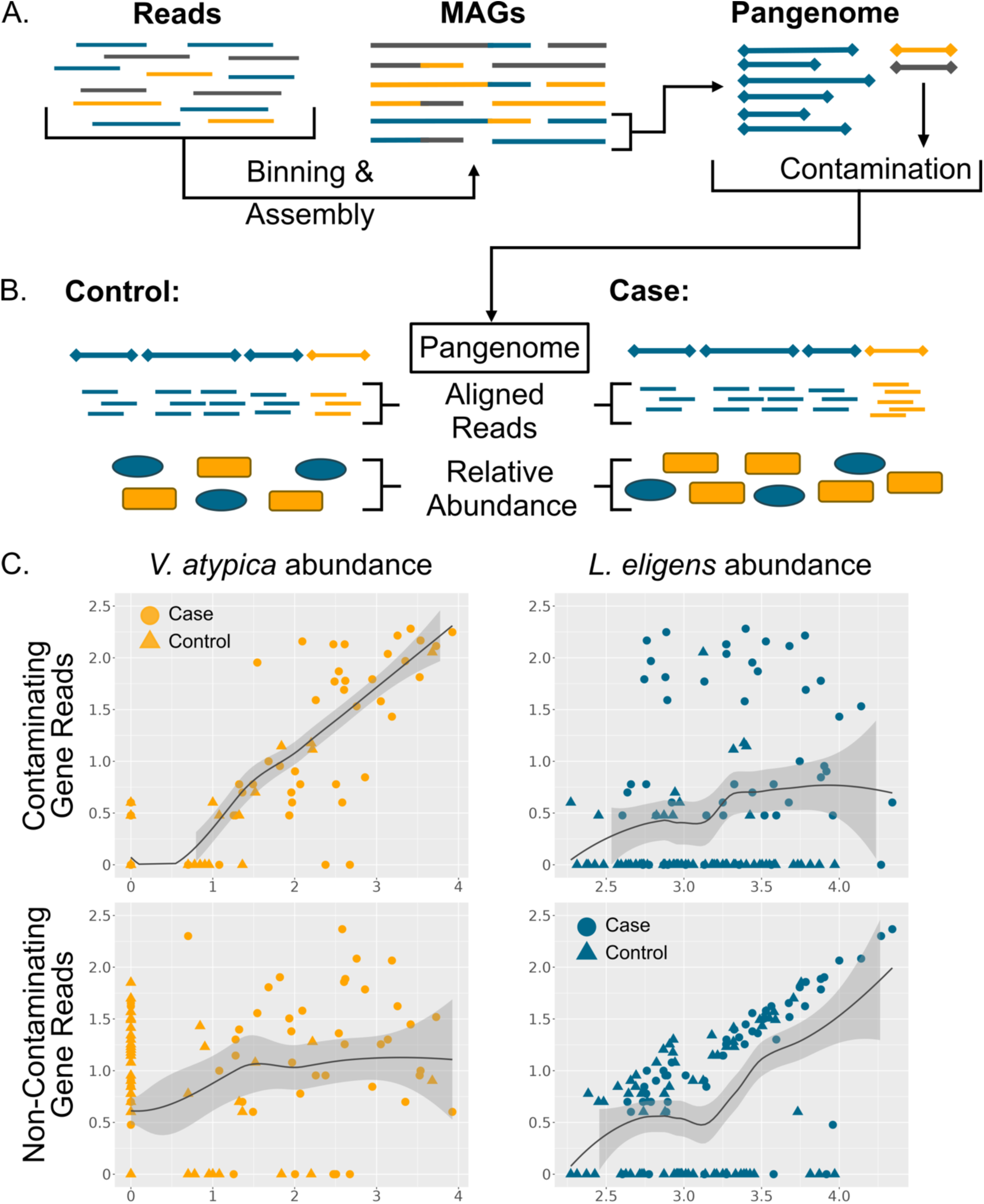
(A) Diagram of the process of assembling shotgun metagenomic reads into metagenomic assembled genomes (MAGs). The true origin of each piece of DNA is represented by color. Here, contamination from the yellow and blue species is introduced into the pangenome of the grey species. (B) Illustration of aligning metagenomic reads from controls (left) or cases (right) to a pangenome (grey) with a small amount of contamination (yellow). The relative abundances of the grey and yellow species are visually represented below. (C) Graphs showing log_10_ read counts of the contaminating species, *Vellonella atypica* (left column) or the originating species, *Lachnospira eligens* (right column) plotted against log_10_ read counts of a contaminant gene (top) or a non-contaminant gene (bottom). All counts had one pseudocount added before log-transforming.

It is unknown how tolerant a pangenome-based analysis would be to these false positives. Here, we present a case study of commensal gut Clostridia in cirrhosis cases versus controls, in which we attempt to detect and correct for pangenome contamination.

## Results

### Case study: commensal Clostridial gene content in cirrhosis

Numerous studies have found that gut microbial communities are altered in cirrhosis (19–22). Patients with cirrhosis show consistently lower levels of commensal Clostridia, especially those in the *Lachnospiraceae* family (19, 20), and the degree of *Lachnospiraceae* depletion correlates with severity (20). Notably, the liver and gut microbiome have a close anatomical link, exchanging metabolites via the bile ducts and portal vein. *Lachnospiraceae* produce metabolites such as butyrate that are associated with liver health, as well as with increased barrier function (23), which is diminished in cirrhosis. Finally, experimental mouse models of liver damage also indicate that *Lachnospiraceae* have beneficial effects. An oral gavage with fecal material, which enriched for *Lachnospiraceae* and butyrate in the gut, was able to reduce liver injury (24); in a mouse model of primary sclerosing cholangitis, a disease involving liver damage that can proceed to cirrhosis, reintroduction of *Lachnospiraceae* was able to reduce liver inflammation and fibrotic lesions (25). We were therefore interested in identifying genetic elements that could allow *Lachnospiraceae* to persist in the cirrhotic gut, as this could help drive the selection of novel probiotics.

To identify such genes and genetic elements, we re-analyzed short-read shotgun metagenomes from a large case-control study of liver cirrhosis from multiple causes (26) (Qin et al.; n=123 cases and n=114 controls). We used MIDAS2 (6) to assign reads to the pangenomes of species derived from the Universal Human Gut Genome database, or UHGG (10), which includes both isolate sequences and metagenomic assemblies. Next, we applied Fisher’s exact test to identify genes from *Lachnospiraceae* that were differentially prevalent in case versus control metagenomes, using the DiscreteFDR method to adjust for multiple comparisons (27). Setting a false discovery rate (FDR) cutoff of 5% yielded 86 significant genes in six *Lachnospiraceae* species.

To find potential homologs of these genes, we performed a BLAST search of their translated sequences against NCBI’s “nr” database (28). We were surprised to find that in 33 out of 86 cases, the closest matches were not found in *Lachnospiraceae* or even other Clostridia. Rather, they were found in distantly related taxa such as *Veillonella* (class Negativicutes), *Haemophilus* (class Gammaproteobacteria), and *Streptococcus* (class Bacilli). To confirm these taxonomic distributions more rigorously, we also matched our sequences to fine-grained groups of orthologs from the EggNOG database (29). This was possible because gene sequences from UHGG had already been clustered into protein families at different amino acid identity thresholds in the companion UHGP database (10); these in turn were annotated using several methods including the tool eggNOG-mapper (30). Our results largely matched what we observed using BLAST (Table 1, Supplementary Table 1).

**Table 1.**
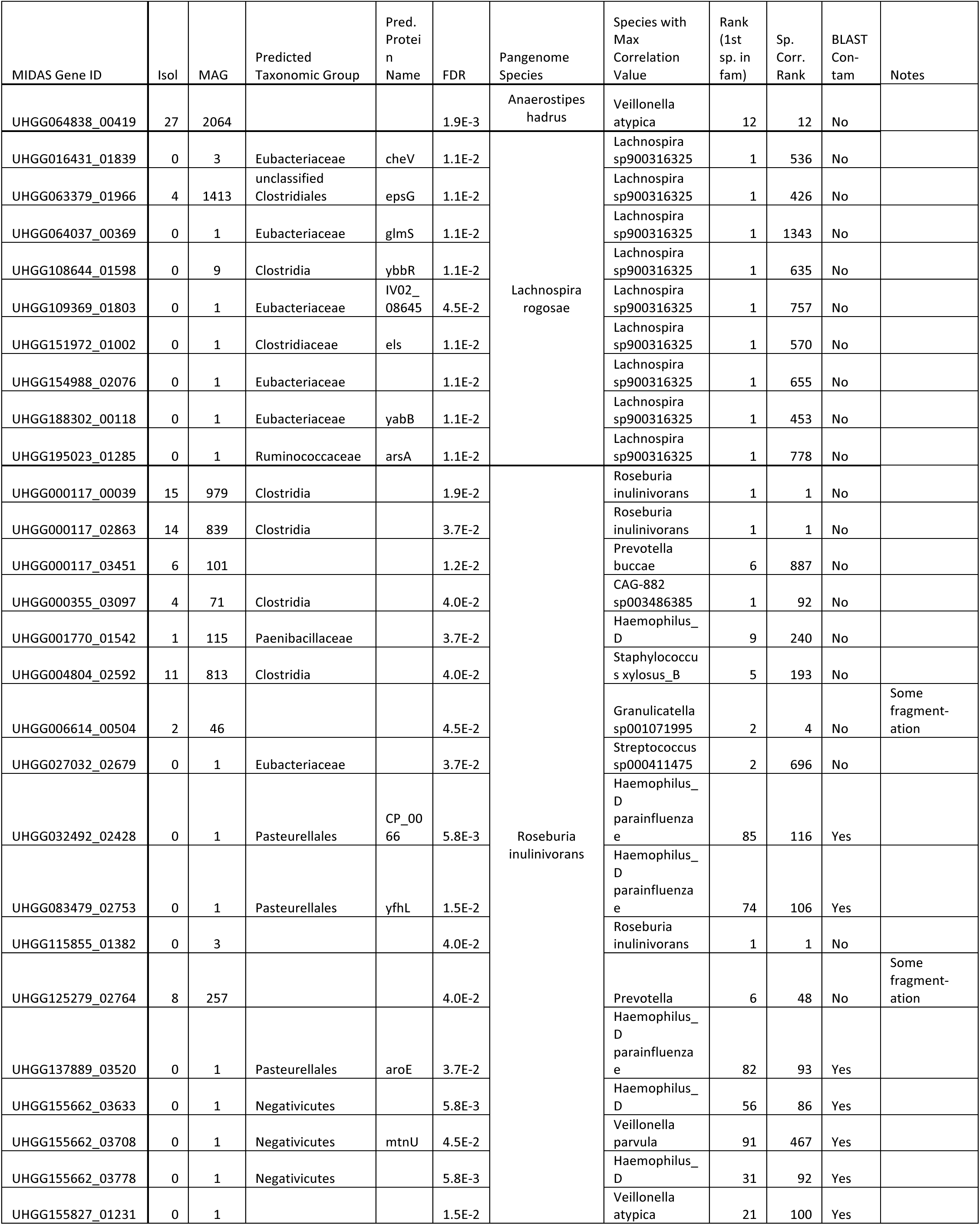
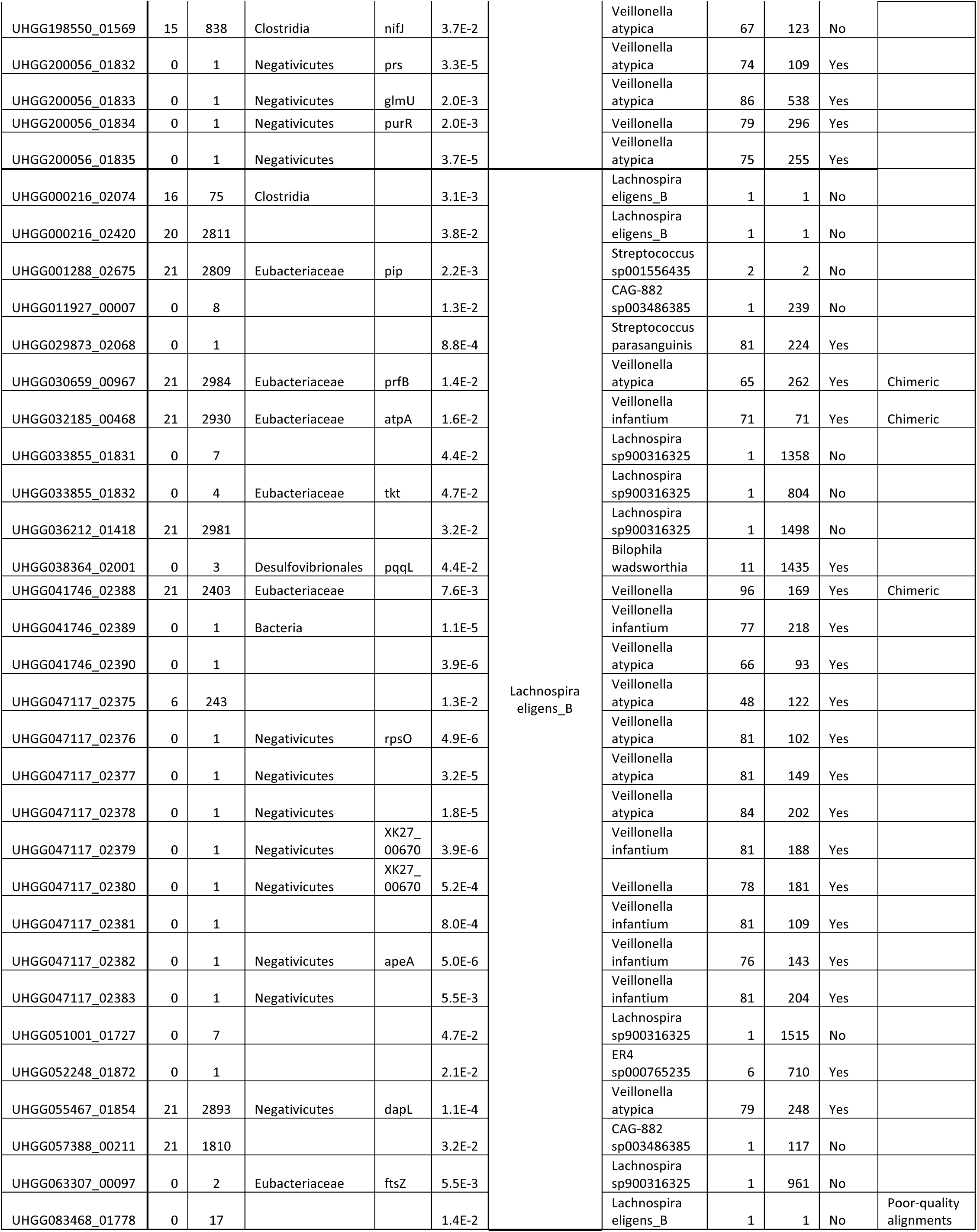
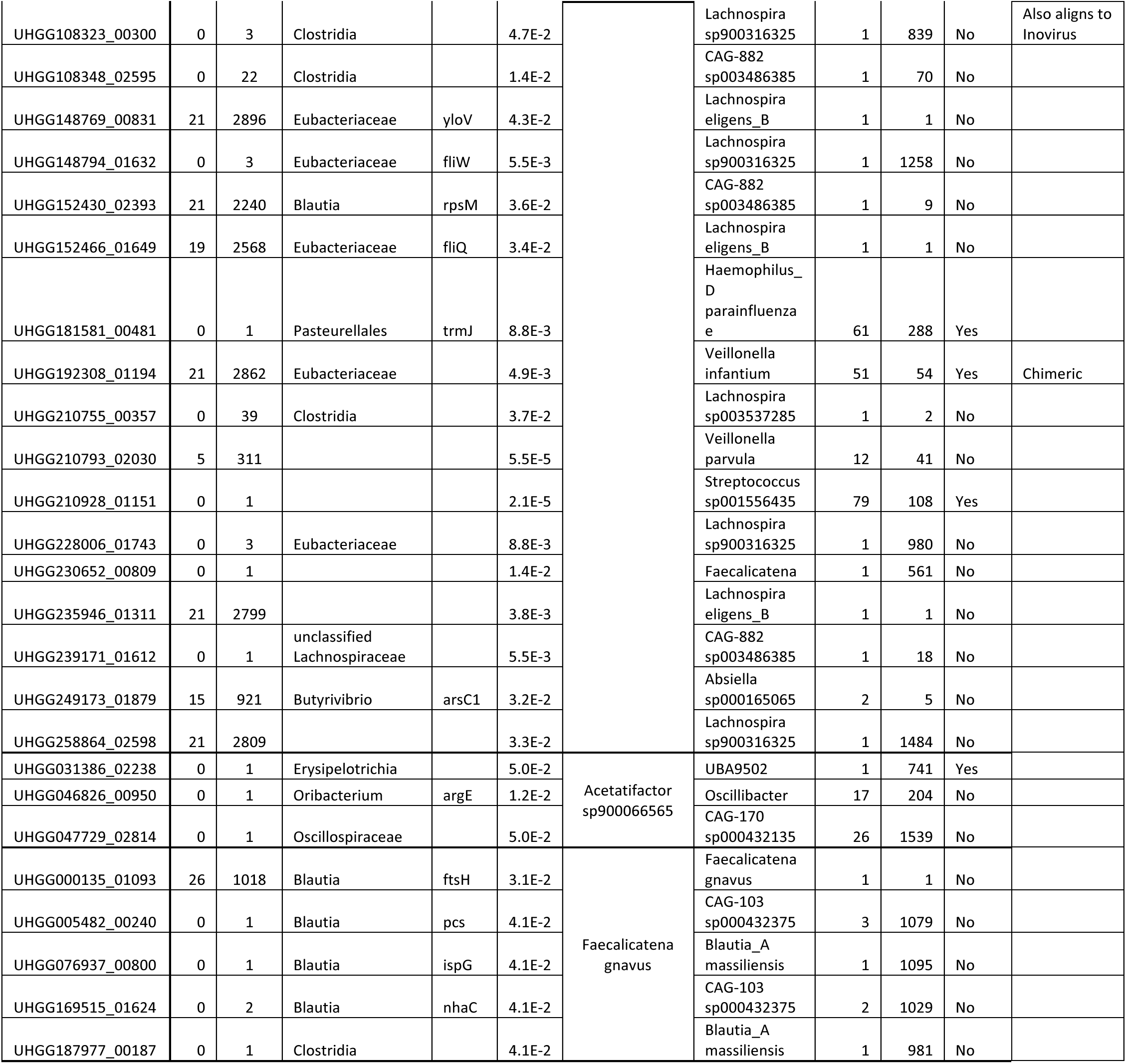
Table showing the PanSweep results for all *Lachnospiraceae* species that had genes significantly associated with cirrhosis. Columns include: 1) the ID of the gene in the MIDAS2 UHGG database; 2-3) the number of solate and MAG genomes in UHGG that contain the gene; 4-5) the predicted taxonomic range and predicted name of the corresponding UHGP-90 protein cluster, using UHGG’s EggNOG-mapper annotations; 6) the false discovery rate *q*-value of the gene; 7) name of the species in the MIDAS2 UHGG database; 8) which species was most correlated to the gene; 9) the highest rank of species in the same family of the originating pangenome; 10-11) whether the BLAST results indicated contamination or not, and any other observations from the BLAST results (e.g., if the gene appeared to be chimeric, or if the alignments were generally poor quality).

Qin et al. (26), as well as other studies (19, 22), found that the oral taxa *Streptococcus, Haemophilus* and especially *Veillonella* were more abundant in cirrhotic gut microbiomes. Thus, the apparent increase in these genes’ copy number could be an artifact caused by increased *Veillonella, Haemophilus,* or *Streptococcus* relative abundance. Indeed, the ABC transporter gene UHGG047117_02377, an example of a likely contaminant, correlates better with the abundance of *Veillonella atypica* than the species of the pangenome where it appears, *Lachnospira eligens*. The opposite trend can be seen for the flagellar gene FliQ (UHGG152466_01649) in *L. eligens* (Figure 1C).

Out of the 33 putative contaminant genes, 26 were present in only a single *Lachnospiraceae* assembly. According to CheckM, these 13 single assemblies had a mean contamination estimate of 0.22% (interquartile range: 0% to 0.24%). This is actually below the average contamination estimate of all *Lachnospiraceae* genomes (1.4%; IQR: 0.19% to 2.2%). Overall, assemblies containing contaminant genes had somewhat higher estimated contamination than other assemblies from the same species, but all were still below the 5% threshold that CheckM recommends (15) (Supplementary Figure 2). A recent revision of UHGG (31) uses an additional, more sensitive chimera detection tool called GUNC (16), but this did not flag any of the assemblies bearing the contaminant genes. These results demonstrate that in pangenome-based analyses, even unremarkable amounts of assembly contamination can lead to pathological results when the contamination source is itself correlated with the phenotype of interest. Furthermore, because misassembly is most likely to result from microbes found in the same environment, this scenario may be more common than it first appears.

### An analysis and visualization tool to identify potential contaminants

To help researchers identify potential contaminants that may obscure pangenome-based analyses, we created a workflow called PanSweep that integrates the tests we performed above and visualizes the results. This workflow, designed to analyze the results from the tool MIDAS2 (6), helps flag contaminants in three ways: (1) loading and comparing metadata from the EggNOG database per gene (29, 30); (2) correlating gene count and species abundance; and (3) visually representing patterns of gene co-occurrence across metagenomic samples.

First, each gene is associated with its UHGP-90 cluster (i.e., clustered at 90% amino acid identity), allowing retrieval of the predicted taxonomic distribution according to EggNOG. This information is reported so that the user can assess whether the lineages match. Second, per-gene read counts are correlated with all species abundances using Spearman’s ρ. For each gene, the correlations to each species are then ranked from highest to lowest. We report the rank of the most-correlated species that matches the gene’s expected lineage (based on the pangenome in which it is found). Finally, to determine which genes may be part of the same contig or mobile element, we calculate a similarity matrix of gene co-occurrence using the Jaccard index. This Jaccard matrix is visually represented as a clustered heatmap and as ordination plots: the user can choose between UMAP (32), NMDS (33, 34), and PCoA.

The gene-to-species correlation test was especially consistent with our BLAST analysis. Out of the 39 genes where the most correlated species was in the same family, only one showed evidence of contamination via BLAST, a false positive rate of 2.6%. This only increased to 2/50, or 4%, when we allowed a family lineage match anywhere in the top 10 most-correlated species. Conversely, in the 28 cases where a lineage match did not occur in the top 50 most-correlated species, 27 were confirmed to be contaminants by BLAST (a true positive rate of 96%). We can summarize the performance across all cutoffs by calculating the area under the receiver-operator characteristic curve (AUROC), where 1 indicates perfect prediction and 0.5 is random. The correlation lineage test had an AUROC of 0.97, indicating near-perfect classification (Supplemental Figure 2). In contrast, a variation where we only considered exact species matches was not effective (AUROC = 0.53, near random).

EggNOG taxonomic annotations were also highly predictive of contamination (AUROC = 0.86). All 19 cases in which EggNOG predicted a protein’s range to be in the Negativicutes or Pasteurellales were confirmed by BLAST analysis to be contaminants, with similar sequences mostly or exclusively outside of the *Lachnospiraceae*. EggNOG annotations to *Eubacteriaceae*, Clostridia (and similar taxa), *Lachnospiraceae*, or *Ruminococcaceae* tended to be legitimate: 28 of these were likely true *Lachnospiraceae* genes, while four appeared to be contaminants. However, EggNOG annotations were not available or very general (“Bacteria”) for 24 cases, eight of which were likely contaminants based on BLAST.

Gene-to-gene correlation can also help identify contaminants. When we calculated Jaccard dissimilarity between all the significant *Lachnospira eligens* genes and examined the co-occurrence heatmap, 13 of the putative contaminants formed a single well-defined cluster. Nine of these were in fact adjacent features on a single contig in the same MAG, GUT_GENOME047117 (Fig 2C). In contrast, 25 non-contaminant genes in *L. eligens* were not correlated with the genes that failed, and not all were correlated with one another (Fig 2C).

**Figure 2.**
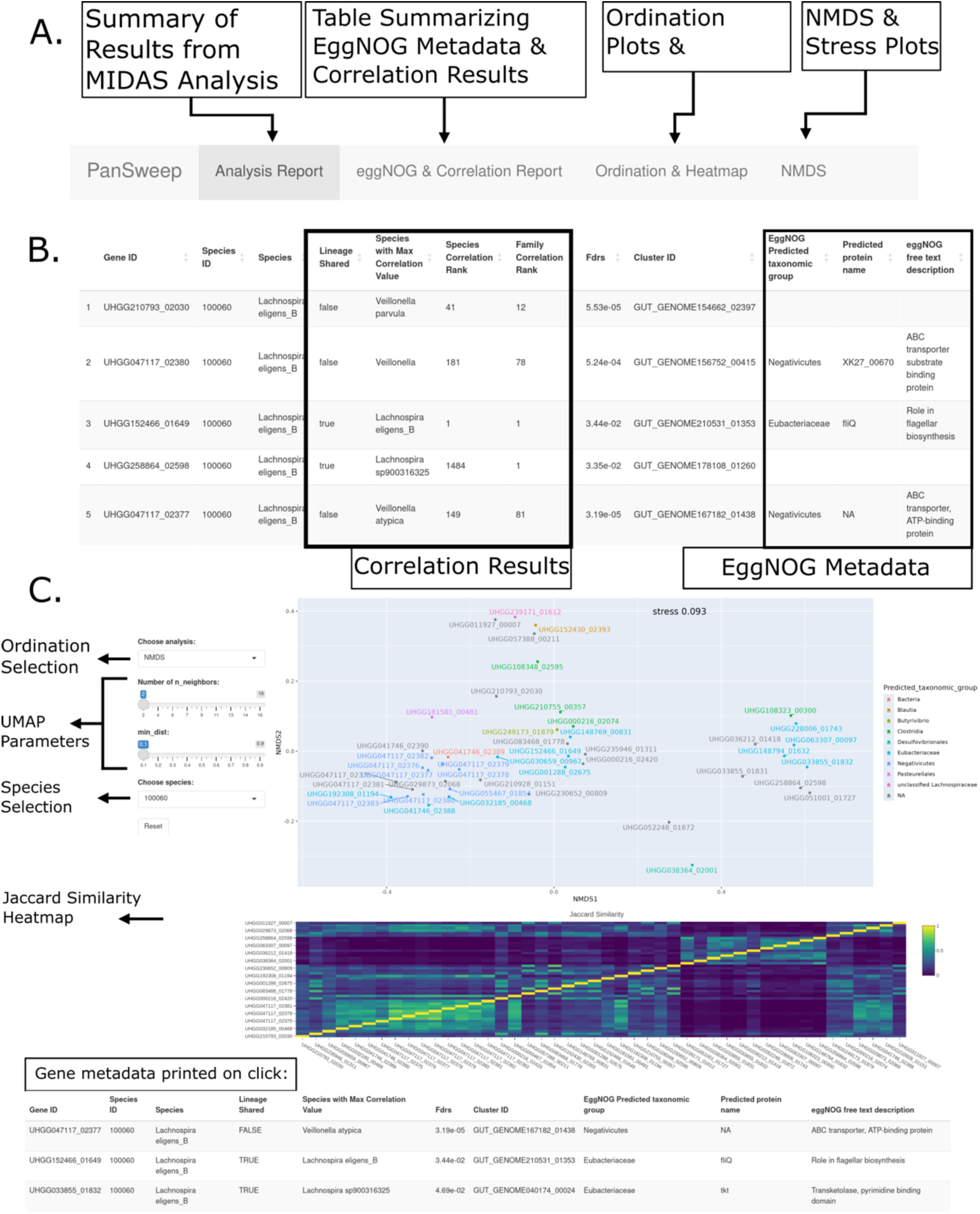
(A) Annotated screenshot explaining the four main tabs of PanSweep. (B) Annotated screenshot showing the first five rows of a table of significant genes, including the lineages from the eggNOG metadata and the correlation test. The columns from left to right show the gene id in the UHGG database, the species id rom MIDAS, the species name, whether the lineage of the most correlated species was shared with the gene’s originating species, which species was most correlated to the gene, the calculated FDR, the UHGP-90 cluster dentity, the predicted taxonomic group from the eggNOG database, and the predicted protein name from eggNOG. (C) Ordination and heatmap for *L. eligens.* For this static visualization, the interactive NMDS plot of gene presence-absence (Jaccard distance) is zoomed in around one cluster, and gene labels have been added o each point. The sliders are provided to allow changing UMAP parameters. Finally, the Jaccard similarity of gene presence-absence profiles across samples is shown as a heatmap below the NMDS.

Overall, 38% (33/86) of our significant results were likely to be the result of contamination. Importantly, statistical significance did not predict contamination, and in fact, was anti-correlated (AUROC = 0.21), meaning that stricter cutoffs would actually make the problem worse. Considering just those genes with adjusted p-values of below 0.01, 72% were contaminants (23/32).

### Removing contaminants enhances biological signal

The majority of the 25 *L. eligens* non-contaminant genes had no predicted protein name or description from EggNOG, highlighting the need for improved functional annotation of gut microbial genomes (10). Strikingly, however, out of the nine genes that had any EggNOG free-text annotation, two were directly involved in chemotaxis, and two more had a plausible role in this process. A gene annotated as FliQ, part of the type III secretion system that helps to assemble the flagellar hook-basal body (35), was found in 80% of cases (41/51) but only 48% of controls (37/77). In contrast, a gene annotated as the flagellar regulator FliW had the opposite trend, being found in 26% of controls (20/77) and zero cases (0/51). An unnamed third gene, UHGG228006_01743, had a similar pattern to FliW (19/77 controls, 0/51 cases). Searching this third gene’s amino acid sequence in InterPro (36) revealed the presence of a GGDEF diguanylate cyclase domain (37) and a PAS sensory domain (38). (While the EggNOG annotation also suggested the presence of an EAL domain, we did not find any evidence for this in InterPro.) This combination of domains suggests the possibility that in response to the presence or absence of an intracellular signal, the protein UHGG228006_01743 may synthesize cyclic di-GMP, a metabolite that typically regulates the transition between motility and other behaviors like biofilm formation (39). Another unnamed diguanylate cyclase was also differentially prevalent but was more frequently seen in cases (1/77 controls vs. 13/51 cases).

To investigate why we saw both positive and negative associations with cirrhosis, we plotted the estimated copy numbers of all genes in the *L. eligens* pangenome described in EggNOG annotations as flagellar, plus the two diguanylate cyclases named above (Supplementary Figure 3). This analysis revealed large, distinct clusters of correlated flagellar genes, which we expected since many of these genes are located in the same genomic neighborhoods. One of these clusters included FliQ as well as nine other genes, and had copy numbers that were higher in cirrhosis (adjusted p=0.027, see Methods). A separate cluster included the FliW we identified as significant, as well as the unnamed GGDEF-PAS gene and three other genes, and was strongly enriched in controls (adjusted p=1.1×10^-6^, see Methods). This analysis shows that strain variation in *L. eligens* flagellar loci can differentiate cirrhosis cases and controls. Furthermore, it illustrates that by eliminating likely contaminants, functional linkages – in this case, between *Lachnospiraceae* motility and the cirrhotic gut – can become clearer and more consistent.

## Discussion

This study illustrates an issue with pangenome-based analyses of metagenomes: even small amounts of pangenome contamination can cause the results to be dominated by contaminants. The above case study demonstrates that this problem is not merely theoretical, but occurs even in non-pathological, real-world datasets. This problem is also distinct from another important, previously raised issue called cross-mapping, in which reads do not map uniquely to a single gene (40). In our case, it seems the contaminant genes are being measured accurately, as they correlate best to their likely source. The problem is that trace pangenome contamination leads to copy number variants being attributed to the wrong species.

In this study, the most common sources of contaminant genes were from the *Veillonella* and *Staphylococcus* genera. *Staphylococcus* and *Veillonella* increase in relative abundance in cirrhosis cases, and in fact, increased oral taxa correlate with disease and dysbiosis more broadly (41). A recent explanation for this phenomenon is that oral taxa are actually a biomarker of lower total gut biomass, as they will appear to increase in relative abundance when total gut biomass drops (42). Contamination from these species may therefore be especially problematic for studies of other diseases and the gut microbiome. Strikingly, *Veillonella* was also the source of contamination in a case study (14) used to illustrate that the marker-gene approach used in CheckM (15) failed to identify significant chimerism in an oral *Saccharibacteria* MAG.

The contamination we observed originated mainly from single MAGs, suggesting that dropping infrequently observed gene clusters from the pangenome, an alternative strategy that can be used in the forthcoming MIDAS3 (43), could also be a helpful approach. However, while thousands of MAGs exist for some genera, only a handful may exist for others. Furthermore, in the species *Lachnospira rogosae,* six out of nine *non*-contaminant genes were only observed in a single MAG; dropping these would therefore decrease sensitivity. StrainPGC, a strain deconvolution tool that uses the output of MIDAS3 (43), also uses correlation to refine its estimates of strain genome content. The current study reinforces this strategy, and shows it may be productive to apply a similar technique at the species as well as the strain level.

While our work focuses on a particular case study in the human gut microbiome, we expect that this problem will also be an issue in other environments, especially as more and more metagenomic assemblies are generated. In fact, many environments have markedly fewer available isolate genomes than the human gut, and thus depend more on assembly. Certain genomic features may also pose problems: for example, mobile elements like prophage, which can contain repetitive sequences and/or be present in multiple copies, are known to complicate metagenomic assembly (44).

Fortunately, the method we propose of correlating species and gene abundances appears to be effective, is faster at identifying contaminants than alignments against large databases (yet still agrees with the results), and is broadly applicable across different environments and study designs. This work complements ongoing efforts to decontaminate MAGs themselves by developing and, critically, integrating new methods into MAG assembly pipelines. Methods like MAGpurify (which was applied to some, but not all, of the genomes in UHGG) (45) and MDMcleaner (46), as well as the recently-published deep-learning tool Deepurify (47) and the fast hash-based method FCS-GX (48), will likely be key to constructing more accurate pangenomes as MAG databases continue to expand. As this study demonstrates, removing pangenome contaminants can allow researchers to home in on new associations between microbial gene function and disease, a critical step in moving from species-level associations to testable hypotheses.

## Materials and Methods

### MIDAS2 Analysis

Sequencing runs associated with BioProject ID PRJEB6337 from NCBI’s Sequence Read Archive (49) were downloaded and converted to FASTQ files. We used BBDuk to trim reads from the right below a Phred score of 20, discarding reads that were trimmed below 60bp (50). Samples with fewer than 1×10^6^ reads after trimming and filtering were discarded (n=22), leaving 292 samples. These samples represented 237 subjects, with 123 cases and 114 controls. Trimmed FASTQ files were analyzed using MIDAS2 (6) with the UHGG database (10). Briefly, MIDAS2 was first used to compute species coverage levels across all samples. As recommended, all species with at least 2X marker coverage and 50% uniquely mapped markers in some sample were then selected for copy number variant quantification with MIDAS2. Note that because of this coverage threshold, the total number of subjects in each group differed depending on the species. Copy number estimates for all species in the *Lachnospiraceae* were retained for subsequent analysis. The steps above were performed using a Snakemake pipeline (51).

### Determination of differentially prevalent genes

To identify genes for further study, we focused on prevalence, or the rate that a gene was detected. We used presence/absence estimates generated by MIDAS2. Genes with significant differences in prevalence across samples were identified with Fisher’s exact test, and the resulting p-values were adjusted for multiple comparisons using the Benjamini-Hochberg method (52). Adjusted p-values at or below 0.05 were counted as significant.

### BLAST Analysis

Nucleotide sequences for each gene were obtained from the MIDAS2 pangenomes, which had been clustered at 95% nucleotide ID. Each sequence was first analyzed using NCBI Megablast, with default parameters, against the nt database (53). If no sequences were identified, or the results were ambiguous, the sequence was reanalyzed first using discontiguous Megablast, BLASTN, and finally using translated BLASTX against the nr database. We then used the reported taxonomy information to determine whether most of the best hits were annotated with species falling within the *Lachnospiraceae* family.

### PanSweep pipeline

The PanSweep pipeline is written in R (54) and consists of two parts: a function that performs offline analysis and generates files for visualization, and a function that allows for real-time visualization of the results.

The offline analysis function has three components:

1. Identifying differentially prevalent genes;
2. Calculating Jaccard (dis)similarity of gene presence-absence profiles and performing dimension reduction; and
3. Performing the correlation lineage test.

Part 1 identifies differentially prevalent genes between cases and controls based on the output of MIDAS2 and provided sample metadata as described above. In Part 2, for species with 3 or more significant genes, Jaccard similarities between gene presence-absence profiles are pre-calculated for each gene pair. Then, the Jaccard similarity matrices are converted to dissimilarity matrices, and NMDS, UMAP and PCoA are used to provide two-dimensional representations (55, 56).

In Part 3, for each significantly-associated gene, the gene counts (number of reads mapping to the gene) are correlated to the relative abundance of each species using Spearman’s ρ. We then rank these species-to-gene correlations in descending order, such that a rank of 1 is the highest. Next, the lineage of each species is compared to the lineage of the species where the gene was annotated. We report the highest rank where the lineages match. We consider two versions of this test: one where lineages are matched at the family level, and one where they are matched at the species level.

The real-time visualization tool is a Shiny (57) application that loads in metadata about each gene, plots the above gene-to-gene similarity matrices and ordinations, and displays summary tables showing EggNOG-derived phylogenetic range predictions as well as the results of the lineage test. Jaccard similarity matrices are plotted interactively using plotly (58) with viridis (59) coloring. The user can select genes to highlight on this heatmap; information about selected genes will then appear below. The type of ordination (NMDS, UMAP, or PCoA) can also be set in real-time by the user. Additionally, the UMAP parameters “n_neighbors” and “min_dist” can be user-adjusted via toggle bars, with “n_neighbors” ranging from two to one-third the total number of genes (rounded up to the nearest whole number), and “min_dist” ranging from 0.1 to 0.9 at intervals of 0.1. The stress plot from the NMDS analysis is also provided.

EggNOG predictions were previously made for all UHGP-90 protein clusters (10). To allow the application to efficiently find which UHGP-90 cluster a given gene corresponded to, the tab-separated mapping file provided in UHGP was converted into a Parquet database, with gene IDs stored as integers. We similarly constructed a Parquet database from the eggnog-mapper output provided for UHGP-90 protein clusters. These files are provided with the application. The R package “arrow” (60) is used to read and write Parquet-format files.

### Evaluation

EggNOG taxonomic ranges were calculated using the NCBI taxonomy database, and therefore could not be directly compared to species annotations from UHGG, which uses the GTDB database. We considered annotations to taxa that did not include any *Lachnospiraceae* (*Paenibacillaceae*, Pasteurellales, Negativicutes, Desulfovibrionales, *Erysipelotrichia*, and *Oscillospiraceae*) to indicate contamination. We considered annotations to Clostridia, Clostridiales, *Clostridiaceae*, *Ruminococcaceae*, and *Eubacteriaceae* as consistent with no contamination, because at least some *Lachnospiraceae* have previously been annotated as members of these groups in the NCBI database. When calculating (AU)ROC curves in Supplemental Figure 2, we scored the first group as 1, the second group as 0, and missing annotations or annotations to “Bacteria” as 0.5.

AUROC statistics were calculated using the pROC package in R (61).

### Flagellar copy number analysis

Copy numbers were obtained using MIDAS2. We first obtained the EggNOG-mapper annotations for all genes in the *L. eligens* pangenome (species #100060 in the MIDAS2 UHGG database). We then filtered these annotations for the characters “flag” in the free text description (case-insensitive). We filtered the copy number matrix to include just these genes, plus the GGDEF-PAS domain protein and diguanylate cyclase that we identified as significant in our PanSweep analysis. Descriptive names for each gene were taken from the EggNOG predicted protein name, except in some cases where there was no name or the name only appeared in the free text description, which were added manually.

To test the associations of gene clusters with cirrhosis, we first clustered the 84 genes using partitioning around medoids (PAM) (62) with Pearson correlation distance. We then selected the number of clusters *k* by maximizing the average silhouette width (ASW) over the range *k*=2 to *k*=25, yielding 19 clusters; four clusters had fewer than three genes and were discarded, leaving 15 clusters. For each of these 15 clusters, an eigengene representative was calculated by performing singular value decomposition and taking the first right-singular vector. This eigengene was then tested for an association with cirrhosis using the Wilcoxon rank-sum test; p-values were adjusted using the Benjamini-Hochberg method (52) and a false-discovery rate cutoff of 5% was applied.

Copy numbers were visualized using the ComplexHeatmap package (63), using Pearson correlation to hierarchically cluster genes and Euclidean distance to cluster subjects. We also calculated the quantitative enrichment of each gene for cases or controls using the average *t*-statistic from 500 bootstrap samples of the data.

## Availability

PanSweep is available as an R package at https://github.com/pbradleylab/pansweep under an MIT license. Code to process data and generate figures is available at https://github.com/pbradleylab/cirrhosis-pansweep and at https://github.com/pbradleylab/Updated_PanSweep_Figure_Code. Processed data for reproducing analyses and the Parquet-formatted UHGP databases can be found at https://zenodo.org/uploads/13891285.

## Acknowledgements

The authors thank Dr. Chunyu Zhao and members of the Bradley lab for helpful feedback, and also thank Dr. Zhao for assistance with MIDAS2. This work was funded by startup funds and by NIH grant R35GM151155 to PHB. High-performance compute was provided by the Ohio Supercomputer Center (64).

## Supplementary Information

**Supplementary Table 1:** An expanded version of Table 1 that also includes the UHGP-90 ID each gene maps o, and the total number of genomes it was found in.

**Supplementary Figure 1:** Density plots showing CheckM-estimated percent contamination for genomes containing genes identified as contaminants vs. all other genomes in the same species.

**Supplementary Figure 2:** Receiver-operator characteristic (ROC) curves showing how well the correlation est (“family” and “species”), EggNOG-predicted taxonomic ranges (“EggNOG”), and statistical significance (“FDR”) predict contamination, as ascertained via BLAST. For the correlation test, we report results for the top-ranked match at the family level or below (red, “family”), as well as the top-ranked match at the species level only (purple, “species”).

**Supplementary Figure 3:** Heat map showing MIDAS2-estimated copy numbers of flagellar genes in the *L. eligens* pangenome. Genes are clustered using Pearson correlation and subjects are clustered using Euclidean distance. Genes significantly associated with cirrhosis (Fisher’s test, adjusted p-value ≤ 0.05) are marked in black. The left-side colors show PAM clusters of genes that were significantly associated with cirrhosis (red), controls (blue), or neither (white); clusters with too few genes to perform association tests are colored gray. The right-side “enrichment” colors summarize differences in gene copy number between cases and controls (t-statistic, mean of 500 bootstrap samples; red is enriched in cirrhosis, while blue is enriched in cases).

